# Wound state monitoring by multiplexed, electrochemical, real-time, localized, inflammation-tracking nitric oxide sensor (MERLIN)

**DOI:** 10.1101/2024.11.12.623289

**Authors:** Liyang Wang, Yingqiao Wang, Mabel Bartlett, Daniel San Roman, Gaurav Balakrishnan, Samuel Gershanok, Reem Khan, Clint Skillen, Shanae Butler, Mangesh Kulkarni, Devora Cohen-Karni, Bryan Brown, Tzahi Cohen-Karni

## Abstract

Nitric oxide (NO) released endogenously by induced nitric oxide synthase (iNOS) in macrophages is a key regulatory biomarker for wound inflammation. Detecting NO directly on the wound bed is challenging due to its short half-life time (6 – 50 s), low physiological concentration (nM - µM) and interferences in the complex wound environment. Existing NO sensors suffer from limitations such as rigid substrates, single point of detection and complex measurement setup. Here we present a compliant, multiplexed, electrochemical, real-time, localized, inflammation-tracking nitric oxide sensor (MERLIN) array for in vivo spatiotemporal measurement of NO, with high sensitivity (15.6 ± 5.0 nA/μM for 1.5 mm diameter electrodes), selectivity against nitrites (ca. 27900-fold), ascorbic acid (ca. 3800-fold), and uric acid (ca. 6900-fold), and low limit of detection (8.00 nM). We developed a robust device fabrication protocol, comprehensive and highly reproducible in vitro characterization yielding a consistent device performance deployed for wound healing diagnostics. MERLIN spatiotemporally tracked NO on rat skin wounds for 7 days and results indicated that NO peaks on day 3, in line with previously reported iNOS activity. MERLIN allows spatial mapping of the NO gradient across the wound bed, which can be used to provide diagnostic information to assist wound care.

## Introduction

Nitric oxide (NO), a versatile and ubiquitous bioactive molecule, participates in multiple physiological processes, e.g., neurotransmission,^[1]^ neurovascular coupling,^[2]^ angiogenesis,^[3]^ inflammation and immune response.^[4]^ As a small and uncharged free radical gaseous biomolecule, NO is able to freely diffuse across cell membranes. NO’s extra unpaired electron allows for highly reactive regulatory functions.^[5]^ NO modulates a variety of functions, e.g., activation of soluble guanylate cyclase (sGC) in vasodilation,^[6]^ vesicle exocytosis and neurotransmitter release in neurotransmission,^[1a, 7]^ and phosphorylation of extracellular signal-regulated kinase (ERK) in angiogenesis.^[8]^ In wound healing, proinflammatory cytokine release activates inducible nitric oxide synthase (iNOS) and increases endogenous production of NO,^[9]^ leading to activation of macrophages through S-nitrosylation,^[10]^ and direct DNA damage of pathogens via reactive nitrogen species such as peroxynitrite.^[9b, 11]^ Thus, nitric oxide serves as a biomarker for regulating various physiological processes, including monitoring the progression of the immune response during the inflammation stage of wound healing.^[12]^

Wound healing is a complicated physiological process consists of 4 orchestrated and overlapping stages: hemostasis, inflammation, proliferation and remodeling.^[13]^ The concentration change of signaling molecules, such as cytokines and chemokines, regulates and indicates the transition and completion of wound healing stages.^[14]^ NO has a concentration dependent characteristic which peaks in concentration during inflammation as part of the immune response for antimicrobial effects and decreases when transitioning to the proliferation stage.^[12a]^ During inflammation, iNOS within the infiltrating neutrophils and macrophages at the wound bed catalyze oxidation of L-arginine amino acid (L-arg) to NO with a concentration ranging from nanomolar to micromolar.^[15]^ Measuring NO concentration can provide a quantitative assessment of wound healing process as a point-of-care sensor.

However, *in vivo* detection of NO is challenging due to its low physiological concentration ranging from picomolar to micromolar,^[16]^ short half-life time (6 – 50 s) as a free radical in the biological scavenging environment,^[17]^ short diffusion distance (500 µm)^[17a, 18]^ and the complex bio-environment consisting of ions, metabolic wastes and interferences leading to nonspecific adsorption, and disturbing accurate measurement.^[19]^ To date, various *in vivo* NO detection approaches have been reported including optical resonance,^[20]^ fluorescence imaging,^[21]^ and electrochemical methods.^[22]^ Optical resonance measures NO as a function of refractive index changes and fluorescence imaging technique uses a fluorescent indicator for NO detection. Although these methods reported sub-micromolar NO concentration range, their applicability in real-time *in vivo* NO measurement is restricted by the rigid substrate and complex instrumentation. Electrochemical sensors offer a reliable alternative with advantages of superior temporal resolution, low limit of detection (LOD), and user-friendly setup.^[22]^ Nonetheless, the state-of-the-art electrochemical NO sensor is limited to a single detection point and does not gather sufficient data for physiological process interpretation (Table S1).

The design of an electrochemical sensor for endogenously produced NO requires the consideration of physical properties of NO, i.e., size, charge and hydrophobicity. Redox active NO allows for direct oxidation on the working electrode (e.g., Pt) and the permeable membranes allow selective diffusion by effectively blocking out electrochemical interferences such as nitrites, ascorbic acids, and uric acids.^[17a, 23]^ When NO diffuses across selective layer and reaches the working electrode held at NO oxidation potential (e.g., 0.85V for Pt), the oxidation of NO takes place via a two-step mechanism with an electrochemical reaction (1) which provides electronically readable signals, followed by a chemical reaction (2), described in the following reactions:^[24]^

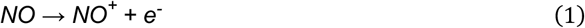

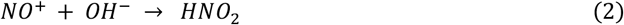

Among various electrochemical techniques, chronoamperometric (CA) detection of NO allows real-time capability with fast sampling rate and high sensitivity.^[25]^ The permselective modification renders specificity towards NO and CA transduces chemical concentration to electrical signals.

Here, we develop a thin and compliant, multiplexed, electrochemical, real-time, localized, inflammation-tracking nitric oxide sensor (MERLIN) array for *in vivo* NO spatiotemporal measurement, with high sensitivity and selectivity. A robust device fabrication protocol was established, along with comprehensive and highly reproducible *in vitro* characterization, confirming consistent device performance (i.e., sensitivity, selectivity, and LOD) for wound healing monitoring. In addition, we used a rodent wound model to demonstrate temporal and spatial NO concentration mapping over the course of wound healing. The recorded NO concentrations matched previously reported literature which informs inflammation trend of wound healing.^[26]^ By using the concentration profile of NO in normal vs. chronic wound healing,^[12b-d]^ MERLIN will enable real-time, reliable and quantitative monitoring of wound state, alleviate subjectivity and dependence of visual expert wound assessment, ^[27]^ thus facilitating chronic wound diagnosis, leading to improved wound care guidance and treatment outcomes.

## Results and Discussion

### Compliant MERLIN array design

*In vivo* NO sensing requires a flexible array that conforms to the wound topography with measurement nodes within NO diffusion distance without impeding wound healing progress.^[17a, 18, 28]^ MERLIN array geometry allows for spatial mapping within a 2-cm diameter rat skin wound model. (Fig. 1a). The scalable, high spatial resolution electrode array (4 by 4 array, electrode diameter 1.5 mm with center-to-center distance of 2 mm) was fabricated following standard microfabrication techniques. Briefly, MERLIN arrays were patterned on a thin polymeric substrate using photolithography, followed by metallization and passivation steps (For fabrication details see Materials and Methods, and Fig. S1, Fig. S2). Finally, to provide a stable electrochemical measurement in the complex wound environment, an on-chip silver/silver chloride (Ag/AgCl) reference electrode was screen printed on the fabricated arrays. The resulting ca. 10-μm thick polymeric substrate and 100-nm thin film metal electrodes are mechanically flexible and can conform to as low as 7mm-radius surface (Fig. 1b), ensuring stable contact with the wound bed.

**Figure 1.**
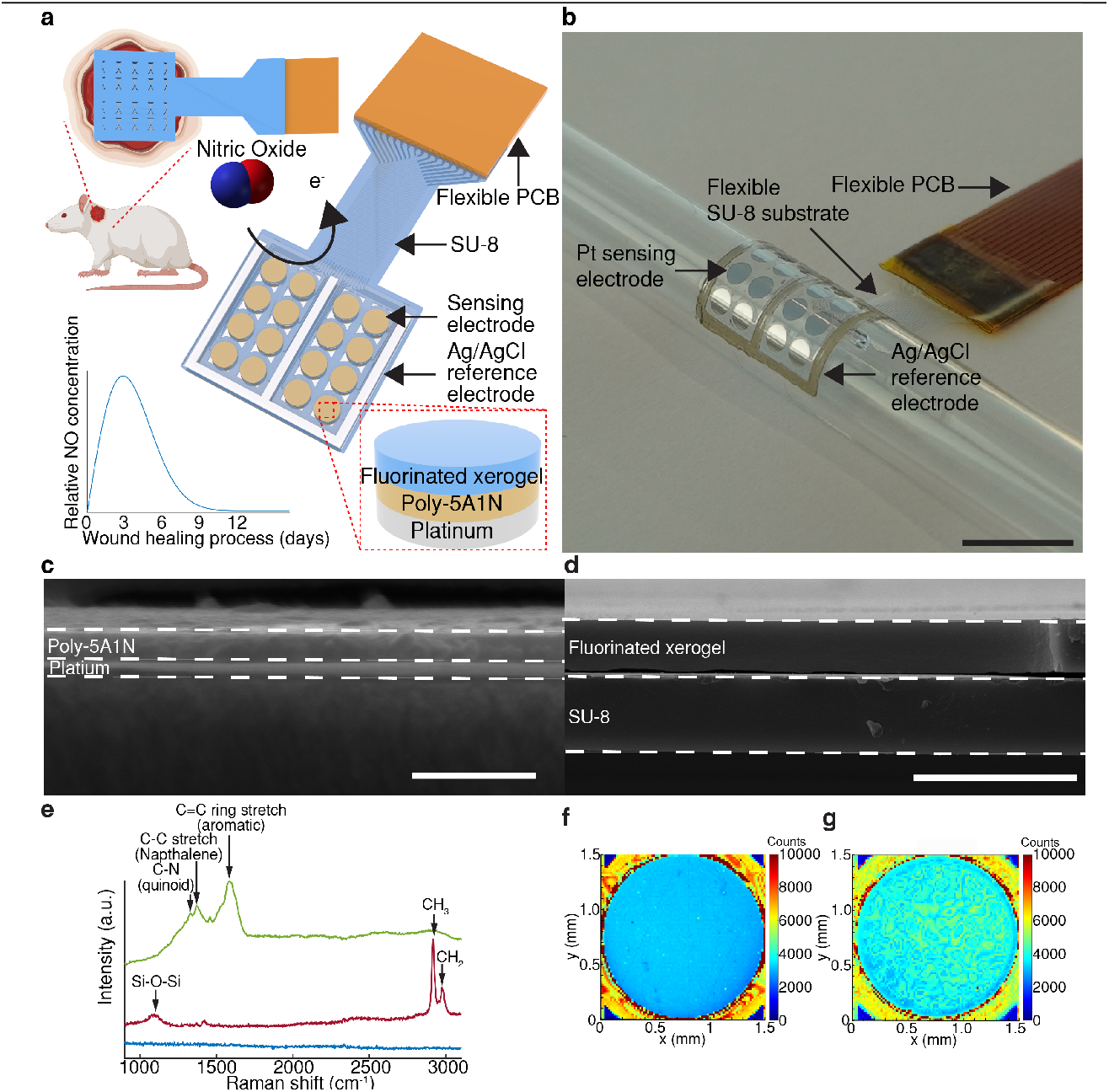
MERLIN for wound state monitoring. **a**, Schematic of MERLIN array. **b**, Photograph of conformable MERLIN array on a 7-mm radius surface. Scale bar, 1cm. **c**, Representative cross-section electron microscopy image of poly(5A1N) on Pt. Scale bar, 500nm. **d**, Representative cross-section electron microscopy image of fluorinated xerogel on SU-8 by spray coating. Scale bar, 10µm. **e**, Representative Raman spectra of modified electrode surface. Blue – Pt; green – 5A1N; red – Fluorinated xerogel. **f**, Representative Raman spectroscopy mapping of 5A1N-Pt electrode plotted at peak intensity of C=C. **g**, Representative Raman spectroscopy mapping of fluorinated xerogel-5A1N-Pt electrode plotted at peak intensity of methyl group.

### Permselective layer for highly sensitive and selective NO measurement

High sensitivity and selectivity towards NO was achieved by permselective layers on the working electrode surface, allowing selective diffusion of NO towards the electrode by effectively blocking out electrochemical interferences such as nitrite, ascorbic acid, and uric acid.^[17a, 23]^ Permselective layers were selected based on size-, charge- and hydrophobicity-exclusion based on NO’s physical characteristics of small size, neutral charge and lipophilicity.^[17a, 23a, 23b, 24a, 29]^ Here, we employed a bi-layer structure of selective materials consisting of electrochemically polymerized 5-amino-1-nathphol (poly-5A1N) (Fig. S3a) and spray coated fluorinated xerogel. The optimization of the deposition parameters resulted in selection of 5 cycles of cyclic voltammetry for fast 5A1N deposition (due to self-terminating characteristics with diminishing improvement in selectivity) and 10 seconds of spray coating to obtain a homogenous layer (Fig. S4).

Electrochemically deposited 5A1N and spray coated fluorinated xerogel exhibited a consistent thickness of 110 ± 3 nm and 2.04 ± 0.17 μm, respectively (Fig. 1c, d, Fig. S3b, for details see Materials and Methods). The presence of poly-5A1N and fluorinated xerogel was confirmed via Raman spectroscopy (Fig. 1e). The presence of 5A1N’s polyaniline-like structures C-N, C-C and C=C ring stretch (quinoid rings, naphthalene, aromatic rings) with peaks at ca. 1330, 1370, and 1590 cm^-1^ respectively,^[30]^ supported its electrochemical polymerization. Crosslinked fluorinated xerogel polymeric coating of the electrode exhibited peaks at ca. 1060, 2840, and 2941 cm^-1^ corresponding to Si-O-Si network, methyl and methylene groups, respectively.^[31]^ Both electrochemically deposited 5A1N and spray-coated fluorinated xerogel layers were uniform as can be seen in the Raman spectroscopy mapping (Fig. 1f, g).

The NO oxidation peak was determined to be 0.82 ± 0.03 V (n = 3) by two-electrode system (Fig.S5a, b, for details see Materials and Methods) which is in good agreement with reported values.^[23b, 32]^ To thoroughly drive the NO oxidation reaction under CA and minimize the reference electrode chemical potential drift, an overpotential above the NO oxidation potential is applied. Thus, the MERLIN is operated at 0.85 V for CA measurement.

To understand the electrochemical properties of the modified electrodes, electrochemical impedance spectroscopy (EIS) was performed. EIS monitors electrode-electrolyte interface properties by applying a small amplitude of alternating potential over a wide range of frequencies and allows the investigation of physical processes at different time scales, e.g., from 500,000 Hz to 1 Hz. The collected EIS data was modelled with equivalent circuit elements which represents electrode-electrolyte interface, including solution resistance (R_s_), bulk capacitance (C_bulk_), double layer capacitance (C_DL_) and electrode resistance (R_electrode_) (Fig. S6).^[33]^ An increase in R_s_ from 797 ± 240 Ω to 5.86 × 10^5^ ± 1.88 × 10^5^ Ω for Pt-poly(5A1N) and Pt-poly(5A1N)-fluorinated xerogel structure, respectively, was observed with deposition of fluorinated xerogel layer which is attributed to decrease in electrode surface accessibility (n ≥ 8, Table S2).^[34]^ Such an increase in solution resistance was also reported in PEDOT:PSS coated nanowire-templated 3-dimensional fuzzy graphene (NT-3DFG).^[35]^ A decrease of C_DL_ from 2.68 × 10^−6^ ± 1.41 × 10^−7^ to 7.32 ×10^−8^ ± 4.32 × 10^−8^ S-s^α^ for Pt and Pt-poly(5A1N)-fluorinated xerogel electrodes, respectively, was attributed to the decrease in electrical conductivity of electrode and increased surface distance from Pt due to sequential addition of selective dielectric layers (Table S2).^[36]^ The decrease in capacitance is beneficial for amperometric sensors through decrease in the non-Faradaic baseline current and increases the signal-to-noise ratio.^[37]^

To ensure the accuracy and stability of on-chip screen-printed Ag/AgCl electrodes, they were compared against commercial Ag/AgCl electrode as the working and reference electrode, respectively, by measuring cyclic voltammetry (CV) in a three-electrode setup in the presence of 1 mM [Fe(CN) 6]^3-^ solution. The CV half-wave potential difference between commercial Ag/AgCl and ink screen-printed Ag/AgCl on-chip electrode is 9.95 ± 0.28 mV (n = 3), which is ca. 1% difference compared to the amperometric operation at 0.85 V (Fig. S7a). The stability of screen-printed Ag/AgCl electrodes shown by a drift of -0.273 ± 0.114 mV/h (n = 3) obtained from a continuous 12-hour open circuit potentiometry test (Fig. S7b,c), confirms steady electrochemical potential of the on-chip reference electrodes in our < 12 hour planned experimental operation both *in vitro* and *in vivo*.

### Sensor performance: sensitivity, selectivity, LOD, and stability

To evaluate the performance of MERLIN array, a standardized NO sensing calibration procedure was developed and deployed in deoxygenated phosphate buffer saline (PBS) solution. Briefly, MERLIN was polarized, and baseline current was determined at 2 hours post polarization. Subsequently, electrochemical interferences were added: nitrite (500 µM), ascorbic acid (100 µM), uric acid (100 µM), and nitric oxide solution at physiologically relevant concentrations ranging from 50 nM to 6 µM (Fig. 2a, Fig. S8a, for details see Materials and Methods). The solution was stirred to circumvent the diffusion of the reactants to the working electrode (Fig. S8b, for details see Materials and Methods).

**Figure 2.**
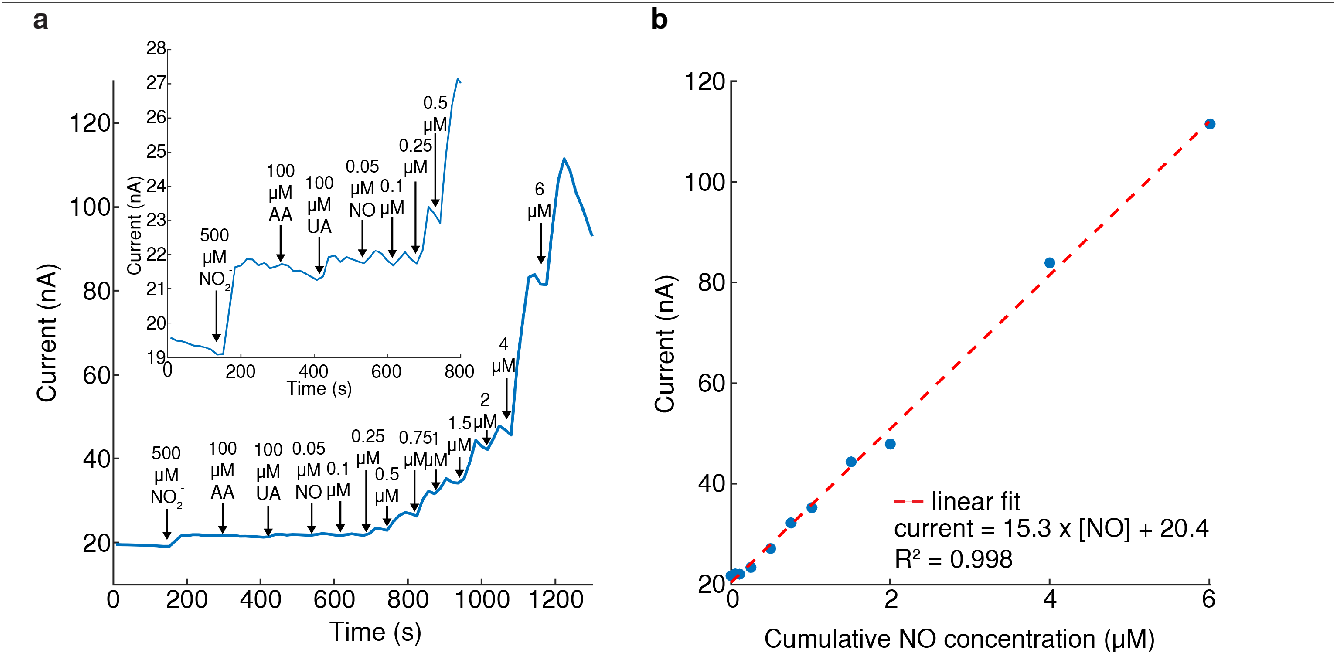
MERLIN in vitro calibration. **a**, Representative single electrode current vs time with NO solution concentration changes. Typical interferants were added, i.e., nitrite (NO$ #), ascorbic acid (AA), uric acid (UA). **b**, Representative calibration curve of current vs NO concentration.

To quantify sensor performance, an electrical current vs. time and NO concentration curves were obtained (Fig. 2). MERLIN demonstrated detection of NO within 2 seconds post analyte solution addition, with a highly sensitive response (15.3 nA/μM) and a linear response up to 6 μM NO (R^2^ = 0.998), within the physiological NO concentration reported in literature.^[16]^ Selectivity against nitrite, ascorbic acid and uric acid was calculated as the logarithmic ratio between sensitivity towards NO divided by sensitivity towards interference (for details see Materials and Methods).

To evaluate a large number (n = 343) of sensors performance for reliable NO sensing experiments *in vivo*, calibrations of MERLIN arrays were implemented with 8 channel multiplexed measurements and were reproducible and consistent (Fig. S9). MERLIN exhibits NO sensitivity of 15.6 ± 5.0 nA/μM (Fig. 3a, n = 343), LOD of 8.00 ± 2.37 nM (Fig. 3b, n = 343), high selectivity against nitrite, ascorbic acid, uric acid at ca. 27900 ± 2700, 3800 ± 500, 6900 ± 800, respectively (Fig. 3c, n = 343). MERLIN demonstrates high sensitivity of 882 nA/µM/cm^2^ which is 26.8-fold higher than reported NO sensor deployed in *in vivo* applications (Table S1).^[38]^ The high selectivity, sensitivity, comparable LOD, multi-channel measurement with high spatial resolution, and extensive sensor calibration with reproducible results of MERLIN, ensure accuracy of NO detection *in vivo* and further clinical applications.

**Figure 3.**
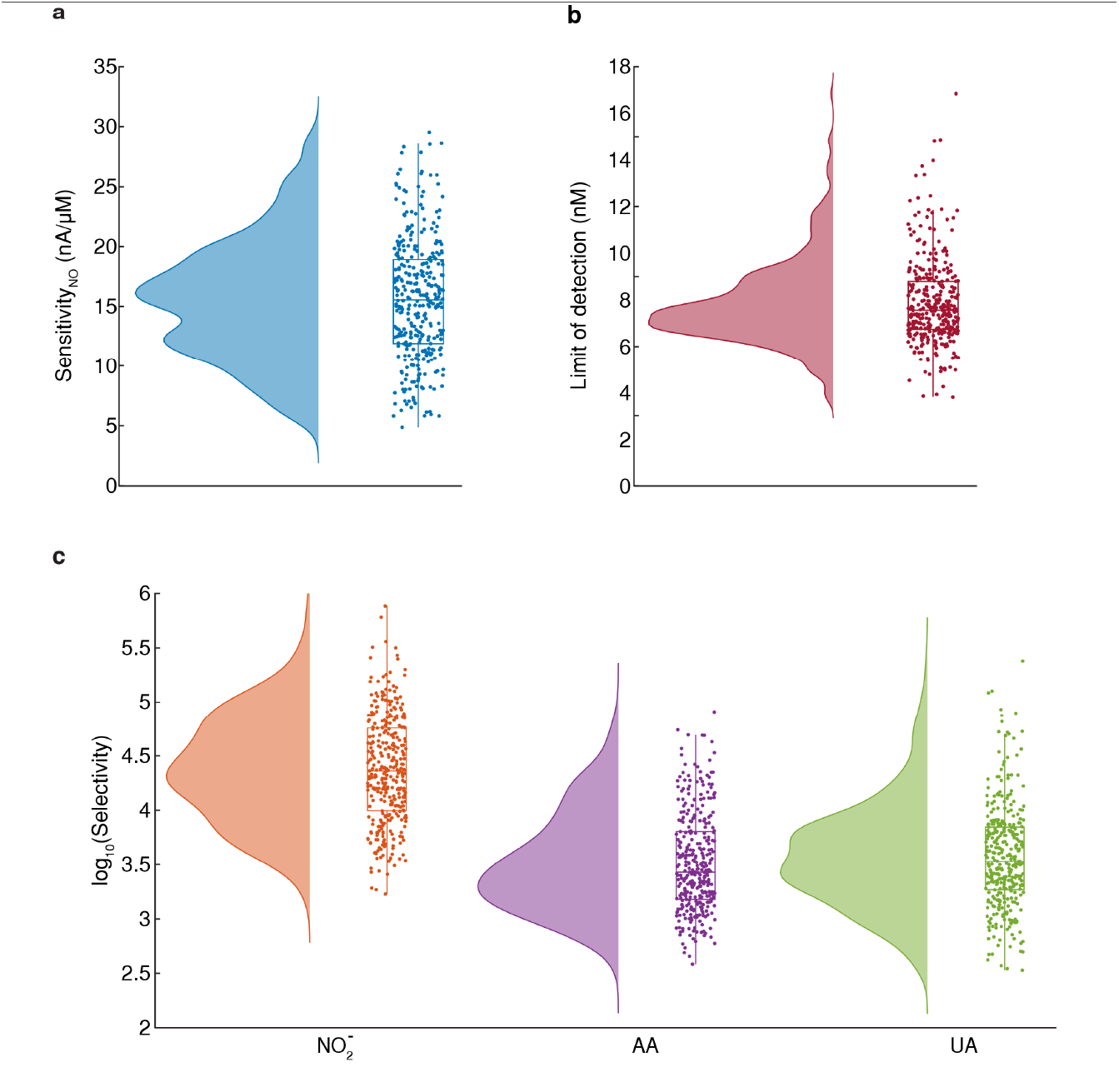
MERLIN in vitro sensing performance. **a**, Sensitivity summary of MERLIN towards NO. (n = 343) b, Limit of detection summary of MERLIN. (n = 343) c, Selectivity summary of MERLIN against common interferences such as nitrite (NO2 -), ascorbic acid (AA) and uric acid (UA). (n = 343); Raincloud plots in **a, b** and **c** include half violin plots showing data distribution, boxplots, and raw data.

### Tracking inflammation during wound healing

To validate *in vivo* NO measurements, acute NO sensing measurements were performed on a rat skin wound model, on day 1, 3, 5, and 7 post-surgery (Fig. 4a). At each timepoint, a freshly calibrated and sterilized MERLIN array was placed directly on the wound to perform 1-hour measurement on anesthetized rats with 8-channel multiplexed measurement (Fig. 4b). NO concentration was extrapolated from converting exponentially fitted baseline current to concentration by sensitivity of each electrode measured through *in vitro* calibration. (Fig. S10a, for details see Materials and Methods).^[34, 39]^ To validate the functionality of MERLIN array, 25 µL of 10 mM L-arg, iNOS substrate,^[40]^ was added 30 minutes post recording to simulate NO production, and was followed by an increase in measured current (Fig. S10b). NO concentrations of 1.77 ± 0.89, 3.55 ± 2.02, 2.55 ± 1.13, 1.63 ± 0.82 μM were detected on days 1, 3, 5, and 7 of wound healing, respectively, using MERLIN. The highest concentration of NO was observed on day 3, indicating a peak in the inflammation phase of wound healing (Fig. 4c). This observation is in line with previous literature on inflammation and NO concentration peak period as demonstrated by iNOS staining.^[12a, 26, 41]^ NO is prominently produced by iNOS, activated by endogenous or exogenous danger signals such as damage-associated molecular pattern (DAMPs) and pathogen-associated molecular patterns (PAMPs) during inflammation stage, with peaks ranging from 0 to 4 days post wound creation.^[13a, 41-42]^ Individual rat NO measurement also indicates an NO concentration peak on day 3 post-wounding, with 3 out of 4 rats (rat 1, 3, 4) confirming this trend, indicating a repeatable and reliable inflammation tracking using MERLIN (Fig. 4c, Fig. S11). Rat 2 was observed to have a delayed NO peak on day 5 which may be due to animal to animal variation in wound healing.^[12a, 15c]^

**Figure 4.**
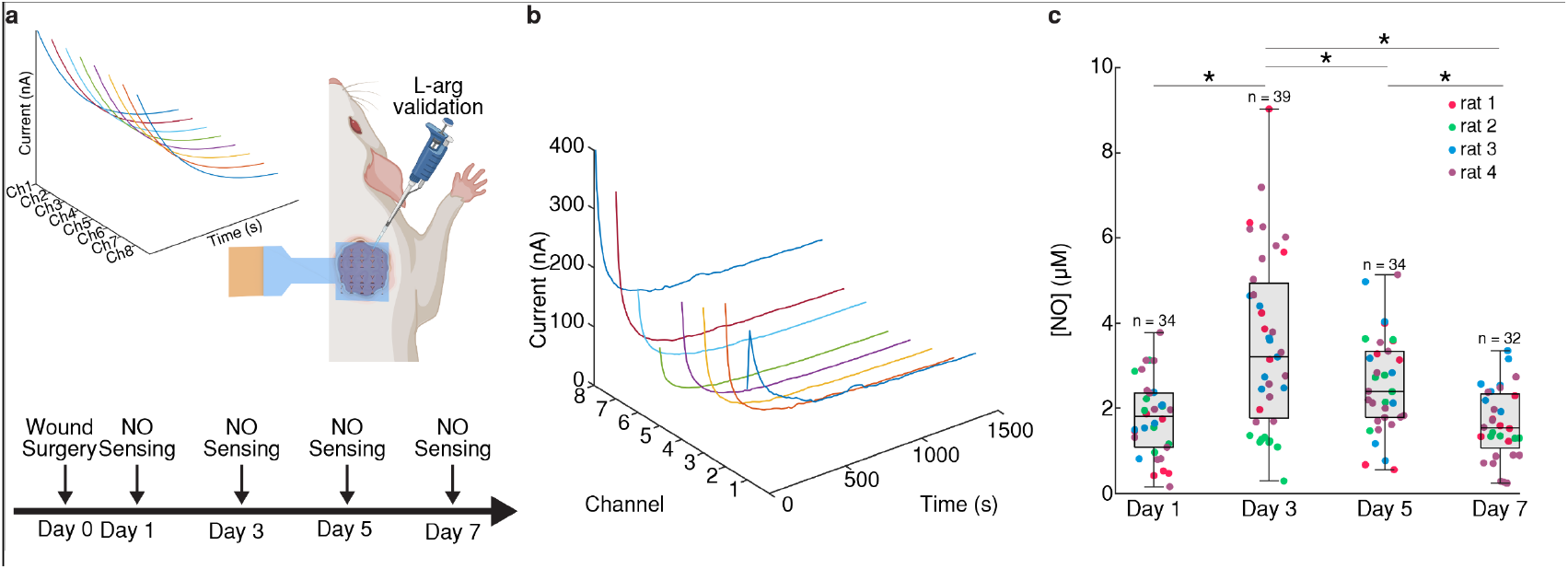
Temporal NO sensing measurement on rat skin wound *in vivo* using MERLIN array. **a**, Schematic illustration of NO sensing experiment *in vivo*, with sensing measurement on day 1, 3, 5, and 7 post-surgery. **b**, Representative 8-channel multiplexed NO sensing *in vivo*. **c**, MERLIN temporal NO sensing averages (n = 4 rats). * denotes p< 0.05, based on one-way ANOVA and *post hoc* Tukey.

### MERLIN can spatially map the wound state

A key advantage of MERLIN array is its ability to spatially map NO concentrations within the wound bed, which may provide information about the distribution of macrophages and iNOS activity (Fig. S12-S16). The results obtained from MERLIN show a distinct NO concentration gradient at different time points. A steep NO concentration gradient was observed on day 3 with sensor readings ranging from 6.20 μM at wound center to 1.67 μM at wound periphery, with standard deviation of 1.85 μM (Fig. 5a,b & Fig. S16b,f). On day 7, a more evenly distributed and lower NO concentration was observed with a smaller range of 1.55 μM at wound peripheral and 0.87 μM at wound center, with a decreased standard deviation of 0.31 μM (Fig. 5c,d & Fig. S16d,h). This decrease concentration gradient was observed on all 4 rats measured. The steeper NO concentration gradient on day 3 suggested that more varied and upregulated iNOS expression during the inflammatory phase of wound healing. The evenly distributed and lower NO concentration on day 7 may indicate the transition from the inflammatory to proliferation stage of wound healing with initiation of wound closure through downregulation of iNOS by anti-inflammatory cytokines.^[12a, 43]^ MERLIN’s ability to spatiotemporally map NO may be beneficial for diagnosing chronic non-healing wounds.

**Figure 5.**
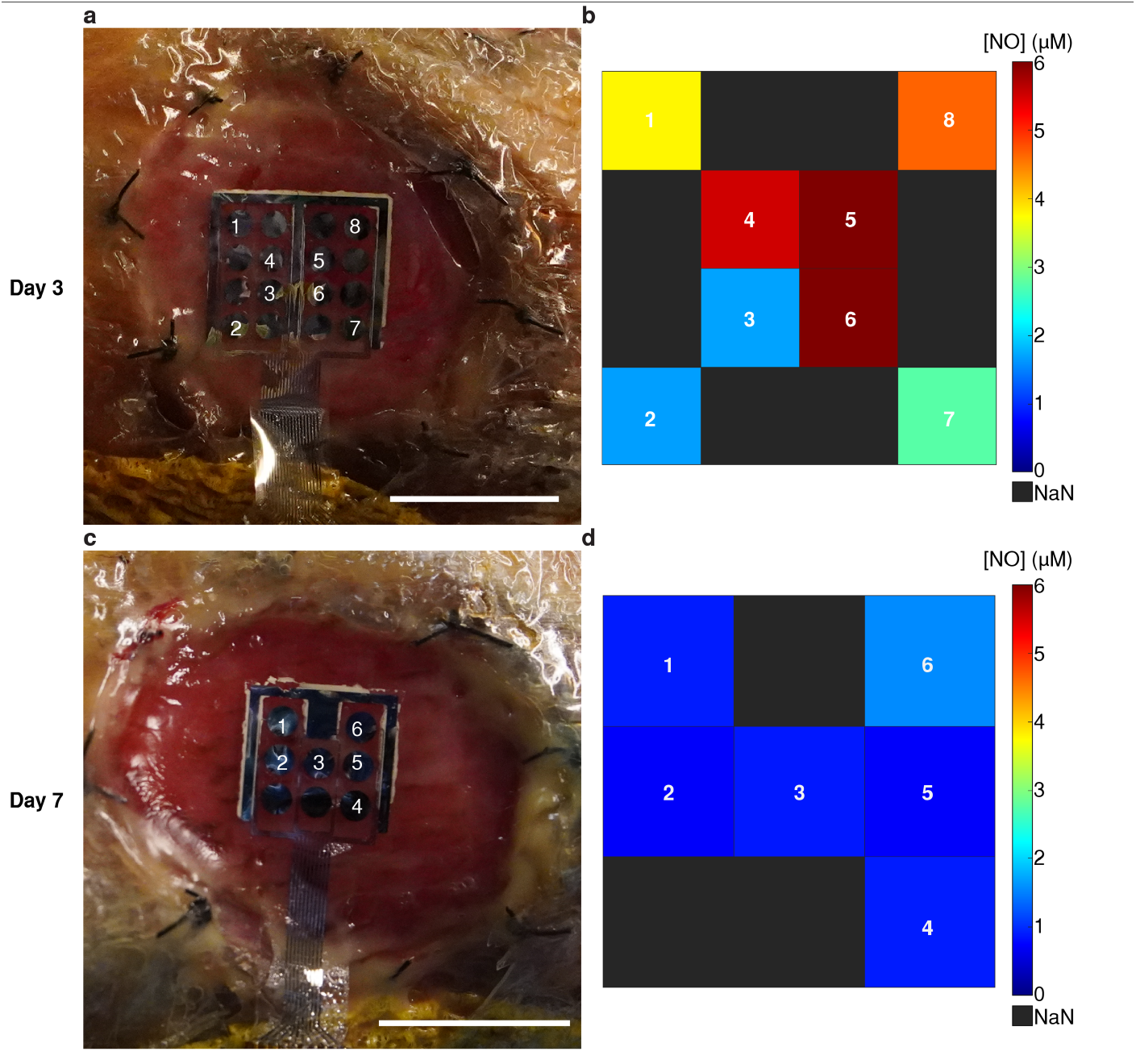
Representative spatial NO Sensing *in vivo* on rat wound using MERLIN. **a**, Representative day 3 MERLIN NO measurement on rat skin wound *in vivo*. Scale bar, 1cm. **b**, NO concentration mapping readout. **c**, Representative day 7 MERLIN NO measurement on rat skin wound *in vivo*. Scale bar, 1cm. **d**, NO concentration mapping readout.

### MERLIN arrays are safe

MERLIN arrays do not elicit adverse tissue response as demonstrated by hematoxylin and eosin (H&E) staining of histological tissue samples (day 7 post wound creation). H&E stained sections showed similar wound morphology of epithelium and granulation tissue (Fig. 6a,b) in NO sensed and in control wounds, suggesting that the sensor placement and measurement did not affect the wound healing process nor did it lead to additional inflammation. Cell density analysis shows a similar number of cell infiltrates observed with 5093 ± 1955 cells/mm^2^ and 5192 ± 1381 cells/mm^2^ in NO sensed and in control samples, respectively, confirming no adverse effects of the NO sensing measurements upon the inflammatory process or wound healing outcome (Fig. 6c).

**Figure 6.**
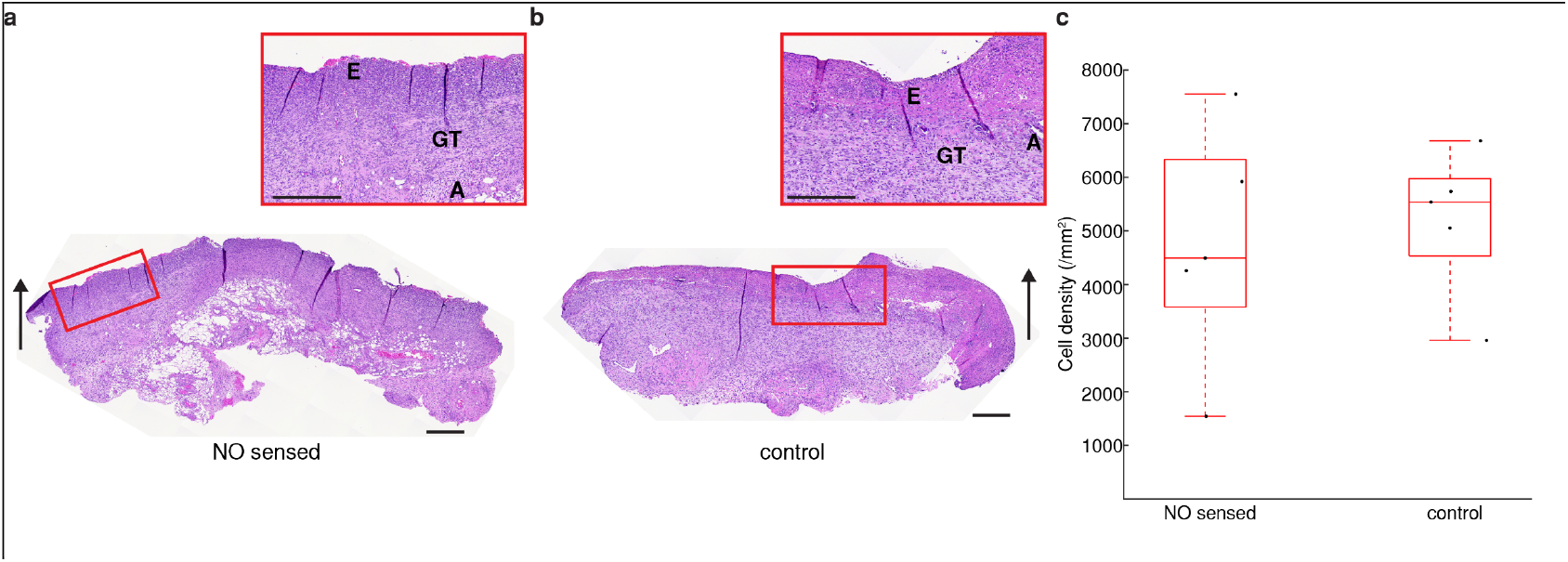
Tissue response to MERLIN. **a**, Representative histological images of tissue harvested from rat skin wound with, and **b**, without MERLIN, at day 7 post-wounding. Both NO sensed wound and control wound show similar wound morphology. Black arrows show the direction of the wound surface. The respective insets show zoomed in regions of the wounds where E = Epithelium, GT = Granulation Tissue, A = Adipose tissue (subcutaneous), scale bar = 400 µm. **c**, Cell density of H&E-stained tissues does not show statistical significance, showing good biocompatibility. (n = 5 rats)

## Conclusion

MERLIN, is a compliant, multiplexed, real-time, highly sensitive and selective nitric oxide sensor array with high sensitivity (883 nA/µM/cm^2^), high selectivity against electrochemical interference (27900-fold against nitrite, 3800-fold against ascorbic acid and 6900-fold against uric acid), spatiotemporal mapping capability, reproducible sensor performance and no adverse effect on tissue and or wound healing. MERLIN provides a practical diagnostic solution for quantitative wound monitoring to complement current practice of visual assessment of wound state. MERLIN can be fabricated as a modular component integrated into a biohybrid electronic platform for feedback regulated wound healing.^[44]^ MERLIN presents breakthrough in quantitative real time measurement of the short-lived NO thus can enable the understanding (and diagnosis) of conditions related to vasodilation, neurovascular coupling, neurotransmission, angiogenesis, and immune response.

## Methods

### MERLIN array microfabrication

MERLIN arrays were fabricated by standard clean room fabrication techniques. Briefly, Si/SiO_2_ (600 nm, Nova Electronic Materials, catalog no. CP02-11208-OX) was cleaned by sonication in acetone for 5 min, rinsed with IPA and N_2_ blow dried. Wafers were treated in barrel etcher (IPC3000 Branson) with oxygen plasma at 850 mTorr, 100 W radio-frequency power for 1 min. The 5 µm thick SU-8 (Kayaku Advanced Materials) was spin coated at 3000 rpm for 40 s prior to 2-step pre-exposure bake at 65 °C and 95 °C respectively for 5 min. UV exposure for photolithography was performed by MA6 (Karl Suss) at 5 mW/cm^2^ for 40 s followed by post-exposure bake at 65 °C for 5 min and 95 °C for 1 min. The micropatterned substrate was then developed in SU-8 developer (Kayaku Advanced Materials) for 5 min prior to hard bakin7g at 190 °C for 30 min. For the metal layer, a bi-layer structure of 300 nm LOR3A (Kayaku Advanced Materials) and 1300 nm Shipley1813 (Kayaku Advanced Materials) was spin coated and baked at 190 °C for 5 min and 115 °C respectively for 1 min. Photolithography was performed by mask aligner MA6 (Karl Suss) prior to development in 2.6% tetramethylammonium hydroxide aqueous solution (CD-26 Developer, Kayaku Advanced Materials) for 60 s. The pattern was deposited with a metal stack of 20 nm Cr and 80 nm Pt by DC sputtering at 50 W in a house-built sputtering system. Lift-off process was performed by using Remover PG (Kayaku Advanced Materials) at 60 °C for 30 min. The patterned metal arrays were passivated with 5 µm SU-8 (Kayaku Advanced Materials), repeating the SU-8 bottom layer process.

### Ag/AgCl on-chip reference electrode screen printing

On chip reference electrode was made by screen printing of Ag/AgCl ink (Creative Materials, catalog no. 126-49) by a micro paint brush (Flat 2, Nicpro), covered by 1-mil PET stencil (McMaster-Carr, catalog no. 8567K12). Curing of Ag/AgCl ink was done at 150°C for 30 min.

### Flexible electrode array backend connection

Flexible PCB was designed to match the backend connection pads of MERLIN array by Eagle (Autodesk) and fabrication of flexible PCB was outsourced to PCBWay. Anisotropic conductive film (ACF) adhesive (3M, catalog no. 7303-5MMX35M) was applied on the back of MERLIN array and pre-heated to 70 °C for tacking. MERLIN array was aligned and bonded to a flexible PCB by manual flip chip die bonder (M9, Laurier) at 150 °C for 30 s.

### Flexible electrode array release from wafer

MERLIN array was released from wafer by etching SiO_2_ in buffered hydrofluoric acid (Thermo Scientific Chemicals, catalog no. 044627.K2) overnight and transferred onto glass slides for further chemical modification on electrodes.

### 5-amino-1-naphthol (5A1N) electrochemical polymerization

Electrochemical polymerization was performed by cyclic voltammetry (CV) by using a PalmSens4 potentiostat (PalmSens BV). CV experiments were performed by using a three-electrode system with platinum electrodes as the working electrode, a Pt wire (CH Instruments, catalog no. CHI115) as the counter electrode and Ag/AgCl electrode (CH Instruments, CH111) as the reference electrode. 5A1N power (TCI, catalog no. A0358) was dissolved in a PBS (VWR, catalog no. 392-0442) adjusted with hydrochloric acid (VWR, catalog no. 20246.298) to pH = 1. CV electrochemical polymerization was performed in 10 mM 5A1N solution and cycled from 0.3 V to 1 V (5 cycles; positive direction initial sweep) at a scan rate of 10 mV/s. Electrodes were rinsed in DI water to remove unbounded monomers.

### Fluorinated xerogel spray coating

A fluorinated sol solution was prepared by adding 7200 µL of ethanol (VWR, catalog no. 85651.320), 1260 µL of Methyltrimethoxysilane (MTMOS) (VWR, catalog no. AAAB23594-AK), 540 µL of (Heptadecafluoro-1,1,2,2-Tetrahydrodecyl) trimethoxysilane (17FTMS) (Gelest, catalog no. SIH5841.5), 1960 µL of deionized (DI) H_2_O (PURELAB Flex, ELGA LabWater) and 120 µL of 0.5 M HCl (VWR, catalog no. 20246.298). The solution was stirred vigorously for 1 hour. Sensors were placed on a hot plate set at 80 °C. The sol solution was spray coated onto electrodes with a gravity-feed airbrush (Iwata, catalog no. N4500) pressurized at 50 psi N_2_ (Matheson gas, catalog no. NI300) for 8 s with a vertical distance of 25 cm. Sensors were removed from hot plates 2 min after spraying and allowed to dry in the air for 48 h before testing.

### Scanning Electron Microscopy (SEM)

SEM images were acquired using a field emission gun SEM (FEI Quanta 600). The accelerating voltage was 10 kV and the working distance was 5 mm. All images were acquired at a high resolution of 2048 × 1768 pixels. No additional conductive coating was applied prior to SEM imaging. The thickness of poly(5A1N) on Pt electrodes was measured by using ImageJ with n = 3 samples.

### Raman spectroscopy

Raman Spectroscopy was performed by using LabRAM Soleil™ Raman Microscope (Horiba Scientific) with 532 nm excitation wavelength at a laser power of 9.2 mW. The spectra were recorded through a 50x objective, 16% neutral density filter, an acquisition time of 30 s, and 600 grating. Raman spectra were acquired from 3 independent samples and 3 randomly distributed spots per sample. Raman mapping was measured with point-by-point scanning made and mosaic image stitching features at a resolution of 100 × 100 points for an area of at least 1.6 mm x 1.6 mm with 4 seconds acquisition at each point. Raman mapping for 5A1N-Pt electrode was computed by MATLAB using RGB color representing relative intensity of C=C bond at 1590 cm^-1^. Raman mapping for fluorinated xerogel-5A1N-Pt electrode was computed by MATLAB using RGB color representing relative intensity of CH_3_ group at 2804 cm^-1^.

### Profilometer thickness measurement

Fluorinated xerogel thickness was measured by a profilometer (Tencor alpha step 200) over a scan distance over 2 mm at a speed of 5 μm/s to measure the step height of fluorinated xerogel film. Step height was acquired from 3 randomly distributed spots per sample and from 3 independent samples per batch of fabrication.

### Electrochemical impedance spectroscopy and equivalent circuit modeling

Electrochemical impedance spectroscopy (EIS) was performed on Gamry R600+ potentiostat (Gamry Instruments) in a three-electrode system with NO sensing electrode, platinum wire and commercial Ag/AgCl electrode as working, counter, and reference electrode respectively. EIS was measured at 0V vs Ag/AgCl reference electrode with a 10 mV alternating current potential from 500,000 Hz to 1Hz in 1xPBS solution. Custom equivalent circuit modeling was fitted on the obtained EIS spectra by using Gamry Echem Analyst software. Models were built by model editor and data was fit by simplex method.

### Cyclic voltammetry and open circuit potentiometry for characterizing screen-printed Ag/AgCl on-chip reference electrode

Cyclic voltammetry was performed on Gamry R600+ potentiostat (Gamry Instruments) in a three-electrode system with gold disc as working electrode, platinum wire as counter electrode, screen-printed Ag/AgCl ink or commercial Ag/AgCl as reference electrode. CV was measured in 1 mM [Fe(CN)_6_]^3-^ in 1 M KCl solution (VWR, catalog no. BDH9258-500G) at a scan rate of 200 mV/s.

Open circuit potentiometry was performed on Gamry R600+ potentiostat (Gamry Instruments) in a two-electrode system with painted Ag/AgCl (Creative Materials, catalog no. 126-49) as working electrode and commercial Ag/AgCl electrode (CH instrument, catalog no. CH111) as reference electrode. OCP was measured for 12 h at a sample period of 1 s.

### Saturated nitric oxide solution preparation

A bubbling system was set up with 2 flasks of 1 M NaOH (VWR, catalog no. 97064-476) DI H_2_O (PURELAB Flex, ELGA LabWater), 1xPBS (0.01 M, pH = 7.4) (VWR, catalog no. 392-0442) and DI H_2_O (PURELAB Flex, ELGA LabWater) in a well-ventilated chemical hood (Fig. S8a). The system was purged with ultra-high purity nitrogen gas (Matheson, catalog no. G1959175) for 1 h to remove oxygen gas. Nitric oxide gas (9.5%, Matheson gas, catalog no. G2659782) was bubbled for 1 h to reach a saturated concentration of 200 μM at 0 °C measured by commercial nitric oxide probe (World Precision Instrument). Saturated NO solutions were freshly made each day.

### Determination of NO oxidation potential by linear scan voltammetry (LSV)

Linear scan voltammetry (LSV) was performed by using PalmSens4 potentiostat (PalmSens BV). LSV experiments were performed by using a two-electrode system with NO sensing electrode as the working electrode and on-chip screen-printed Ag/AgCl (Creative Materials, catalog no. 126-49) as the reference and counter electrode. NO sensing electrode was immersed with 25 µM NO in PBS (0.01 M, pH = 7.4) (VWR, catalog no. 392-0442). LSV peak was fitted by using linear baseline subtraction in the PSTrace software.

### MERLIN array *in vitro* sensing calibration

Sensor calibration was performed in a mechanically stirred, deoxygenated PBS solution. Electrochemical interference solutions of nitrite (Sigma-Aldrich, catalog no. S2252-500G), ascorbic acid (Sigma-Aldrich, catalog no. A4544-25G), and uric acid (Sigma-Aldrich, catalog no. U0881-10G) were added to reach a concentration of 500 μM, 100 μM, and 100 μM respectively, before adding the saturated NO solutions. 10 aliquots of saturated nitric oxide solution were added to reach a cumulative NO concentration 50 nM, 100 nM, 250 nM, 500 nM, 750 nM, 1 μM, 1.5 μM, 2 μM, 4 μM and 6 μM (Fig. S8b). A linear regression of peak current to each aliquot of NO solution against the cumulative nitric oxide concentration was plotted. The slope of the linear regression was the sensitivity of NO sensing electrode. The limit of detection (LOD) is calculated by 3 times standard deviation divided by the sensitivity of the electrode. The selectivity of each electrode is calculated by the logarithmic ratio of sensitivity towards nitric oxide and sensitivity towards electrochemical interferences. Sensitivity, selectivity, and LOD are defined based on the following equations:

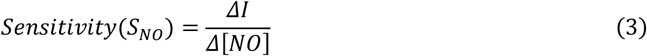

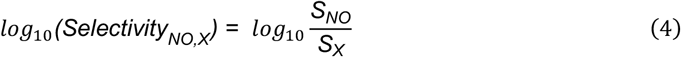

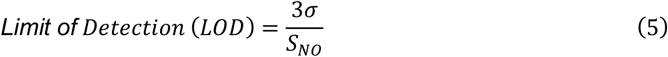

### Rat skin wound model NO sensing *in vivo*

All procedures were approved by the University of Pittsburgh Institutional Animal Care and Use Committee and the DoD Animal Care and Use Review Office (IACUC protocol number 22091435) and were carried out in accordance with the approved guidelines.

Rats were induced and maintained under anesthesia using 2% isoflurane and the surgical site was shaved and prepared in standard surgical fashion. Two 2.0 cm diameter, full-thickness skin wounds were made by sharp incision on either side of the midline in the dorsum of 250-300g male Sprague-Dawley Rats (Charles River, Wilmington, MA). A 2.0 cm inner diameter silicone wound splint (Grace Bio-labs, Bend, OR) was then affixed to the wound margins using 4-0 nylon sutures to prevent wound contraction. The wounds were then covered with a Tegaderm Transparent Film Dressing (3M, catalog no. 7100252805).

NO measurement was performed on day 1, 3, 5, and 7 for a duration of 1 hour on the wound. Sensors were sterilized by 70% ethanol and 30 minutes of UV light. MERLIN arrays were polarized in 1xPBS (VWR, catalog no. 392-0442) for 15 minutes prior to placing on wound. 25 μL of 10 mM L-arginine (Sigma Alrich, catalog no. A5006-100G) was added to stimulate nitric oxide production by iNOS followed by 50 μL of 1xPBS (VWR, catalog no. 392-0442) added as a control.

In a separate cohort of n = 5 rats as the control group, two 2.0 cm diameter, full-thickness skin wounds were made by sharp incision the same way as previously described. Wounds were affixed to the wound margins using 4-0 nylon sutures to prevent wound contraction and were then covered with a Tegaderm Transparent Film Dressing (3M, catalog no. 7100252805). All rats were euthanized on day 7 post-wounding and the wound tissue was excised and fixed in 10% neutral buffered formalin for histologic analysis.

### MERLIN arrays NO measurement *in vivo*

MERLIN arrays were lifted from glass slides by using sharp tweezers. After placing the sensor on the rat skin wound, backend of flexible PCB was connected to a custom break-out PCB board (PCBWay) interfacing with PalmSens 4 (PalmSens BV) connected with MUX8-R2 multiplexer (PalmSens BV) for 8-channel measurement. Sensor array measurement was performed by using method script on PSTrace with 2 s interval per electrode for a continuous measurement of 60 minutes per session.

### Hematoxylin and eosin (H&E) staining

At the time of euthanasia, wound tissue was excised using a 6 mm biopsy punch and fixed in 10% neutral buffered formalin. The tissues were then processed and embedded in paraffin before sectioning at 5 µm and staining with hematoxylin and eosin. Hematoxylin and eosin-stained slides were then imaged at 40X magnification on a whole slide imager (Motic EasyScan, Motic Digital Pathology). QuPath, an open-source image analysis software, was used to quantify cellular density within the H&E images. The number of cells detected within each section was normalized to the area of the section and is reported as cells/mm^2^. Additional qualitative analysis and comparison of wound healing at 7 days in the sensor and control groups was performed by a trained investigator.

## Acknowledgement

T.C.-K. acknowledges funding support from Defense Advanced Research Projects Agency under Award AWD00001593 (416052-5). The content of the information does not necessarily reflect the position or the policy of the Government, and no official endorsement should be inferred. The authors thank Daniel Ranke for providing support for Raman spectroscopy mapping. They also acknowledge support from Materials Characterization Facility at Carnegie Mellon University supported by grant MCF-677785, and Bertucci Nanotechnology Laboratory at Carnegie Mellon University supported by grant BNL-78657879.

## Confiict of Interest

L.W., Y.W., M.B., D.S.R., S.G., and T.C.-K. are inventors on patent applications (PCT/US2024/042777) related to concepts described in this work.

## Data Availability Statement

The data that support the findings of this study are available from the corresponding author upon reasonable request.

