## Supplementary material for "Wound state monitoring by multiplexed, electrochemical, real-time, localized, inflammation-tracking nitric oxide sensor (MERLIN)": Supplemetary information

### Table of contents


| | |
|---|---|
| Notes | Constant phase element modelling to fit for non-ideal double layer capacitance at the electrode-electrolyte interface |
| Figure S1 | MERLIN fabrication process flow |
| Figure S2 | Expanded view of MERLIN |
| Figure S3 | Selective layer deposition and thickness measurement |
| Figure S4 | Optimization of selective layer deposition |
| Figure S5 | Chronoamperometry (CA) operation potential characterization by linear scan voltammetry (LSV) |
| Figure S6 | Equivalent circuit model for electrode-electrolyte interfaces based on EIS data |
| Figure S7 | On-chip reference electrode characterization |
| Figure S8 | MERLIN *in vitro* calibration schematic illustration |
| Figure S9 | Representative 8-channel multiplexed MERLIN calibration |
| Figure S10 | Extrapolation of basal current and MERLIN function validation |
| Figure S11 | Temporal NO concentration summary measured by MERLIN on 4 individual rats |
| Figure S12 | Spatial NO measurement by MERLIN on rat 1 right wound |
| Figure S13 | Spatial NO measurement by MERLIN on rat 2 left wound |
| Figure S14 | Spatial NO measurement by MERLIN on rat 3 right wound |
| Figure S15 | Spatial NO measurement by MERLIN on rat 4 left wound |
| Figure S16 | Spatial NO measurement by MERLIN on rat 4 right wound |
| Table S1 | Comparison among state-of-the-art NO electrochemical sensors to MERLIN |
| Table S2 | Summary of solution resistance, double layer capacitance and chi-square values of equivalent circuit fitting for EIS |
| Table S3 | Calculated constant phase element (CPE) using equivalent circuit models presented in Figure S5 |

### Notes

#### Constant phase element modelling to fit for non-ideal double layer capacitance at the electrode-electrolyte interface

Poly-5A1N modified electrodes and fluorinated xerogel spray-coated electrodes contain an additional capacitive and resistive element related to successive polymeric layer deposition. Due to non-ideal conditions, capacitive elements are modeled as constant phase elements defined by

$$Z_{DL} = \frac{1}{Q_{DL}(i\omega^{\alpha})} \tag{1}$$

where $Z_{DL}$, $Q_{DL}$, $\omega$ and $\alpha$ are constant phase element impedance, constant phase element capacitance, angular frequency of voltage perturbation and constant phase.[1,2]

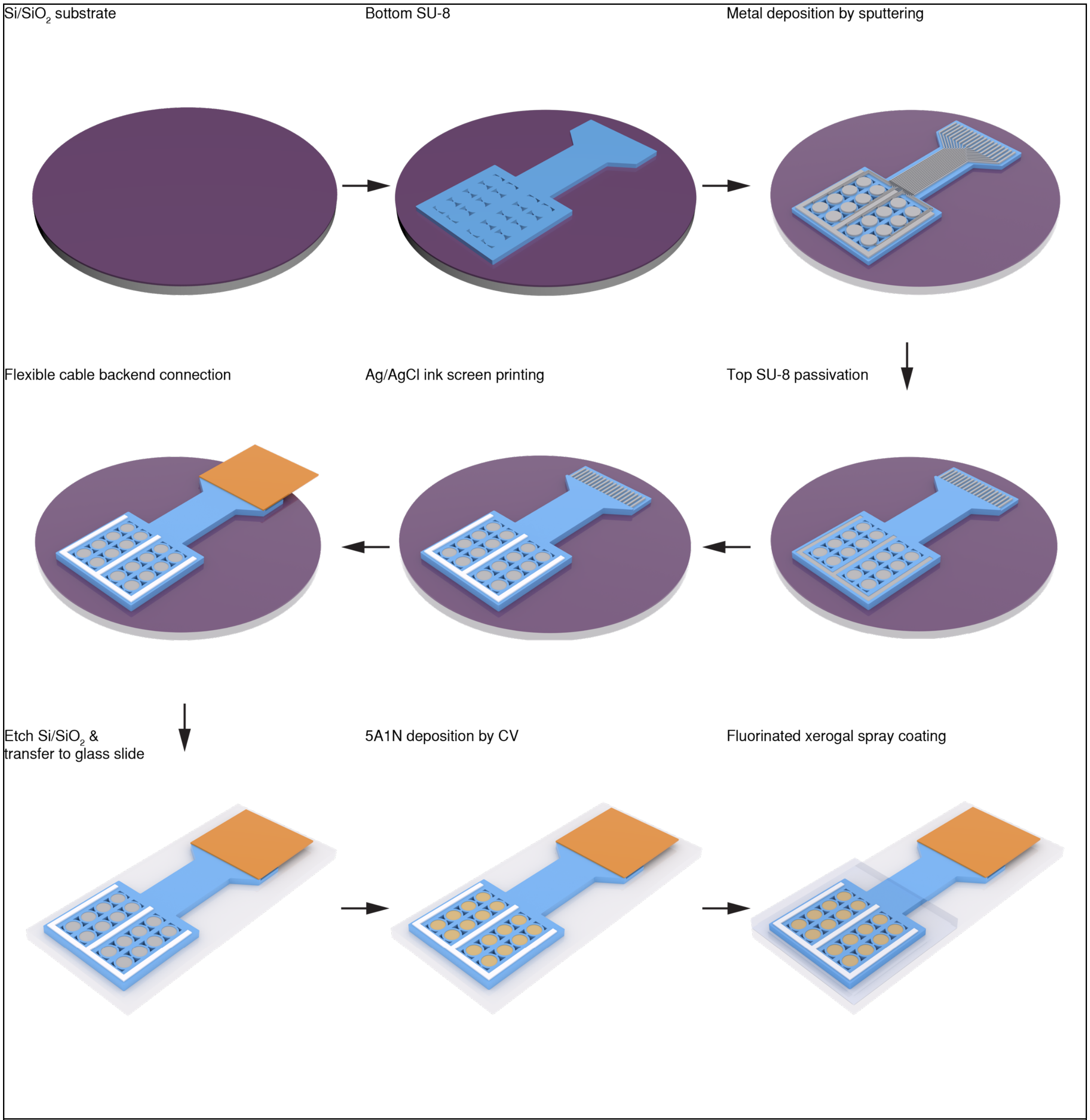


**Figure S1. MERLIN fabrication process flow.** MERLIN arrays were fabricated on Si/600nm $SiO_2$ wafers. 5 μm SU-8 was patterned by spin coating and photobiography. 20 μm Cr and 80 μm Pt was deposited by sputtering, followed by 5 μm SU-8 passivation. Ag/AgCl ink was screen printed as the on-chip reference electrode, followed by flexible PCB connection onto flexible array by ACF adhesives. Flexible arrays were released by etching $SiSiO_2$ in BHF. 5A1N selective layer was deposited by electrochemical polymerization and fluorinated xerogel was deposited by spray coating.

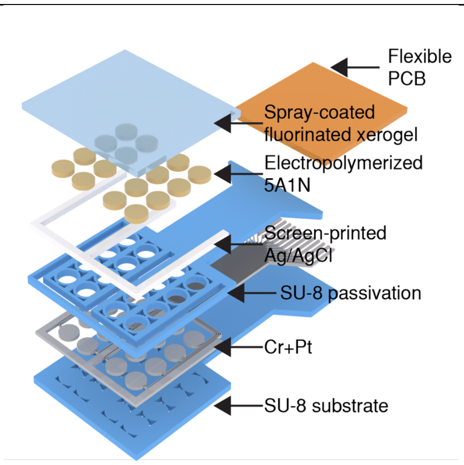


**Figure S2.** Expanded view of MERLIN.

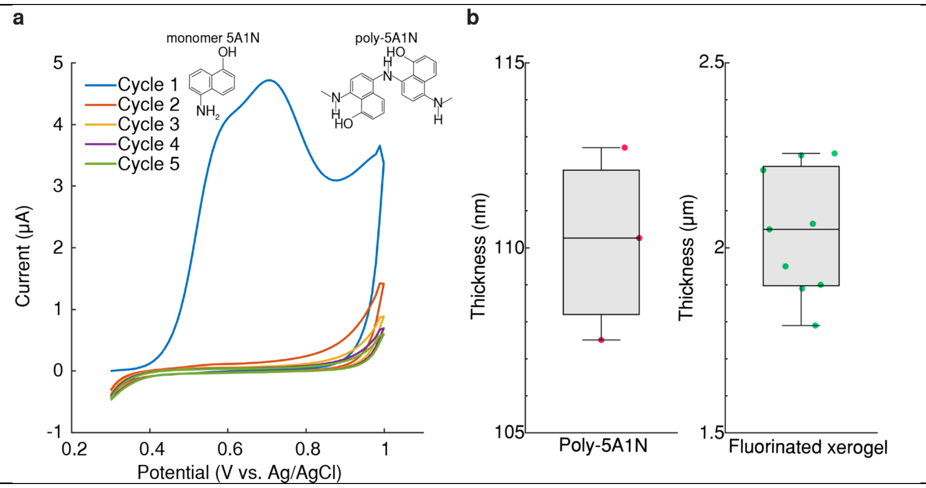


**Figure S3. Selective layer deposition and thickness measurement.** (**a**), Electrochemical polymerization of 5-amino-1-naphthol (5A1N) by cyclic voltammetry. Insert: monomer 5A1N and poly-5A1N. (**b**), Thickness measurement of electrodeposited 5A1N by SEM and fluorinated xerogel by profilometer, n = 3 and 9 for poly-5A1N and fluorinated xerogel respectively.

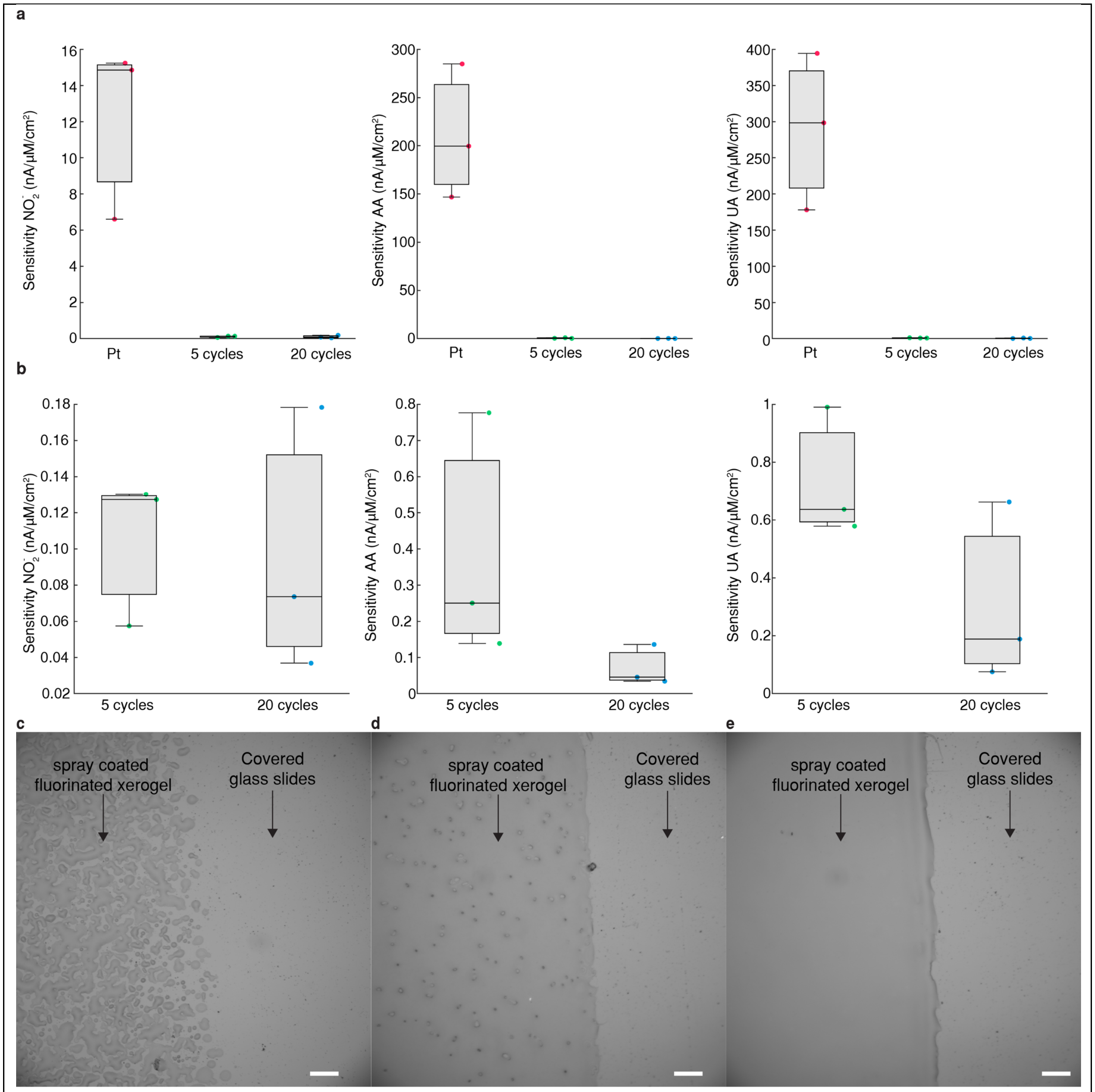


**Figure S4. Optimization of selective layer deposition**. **(a)**, Plot of sensitivity against electrochemical interference such as nitrite, ascorbic acid, and uric acid for plain platinum (Pt), 5 cycles of 5A1N cyclic voltammetry electrochemical polymerization (5 cycles), and 10 cycles of 5A1N cyclic voltammetry electrochemical polymerization (10 cycles). (**b**), Zoomed in plot of sensitivity against electrochemical interference such as nitrite, ascorbic acid, and uric acid for 5 cycles of 5A1N cyclic voltammetry electrochemical polymerization (5 cycles), and 10 cycles of 5A1N cyclic voltammetry electrochemical polymerization (10 cycles). **(c)**, Image of spray coated fluorinated xerogel for 5 seconds, **(d)** for 8 seconds and **(e)** for 10 seconds to achieve a homogenous layer. Scale bar 100 µm.

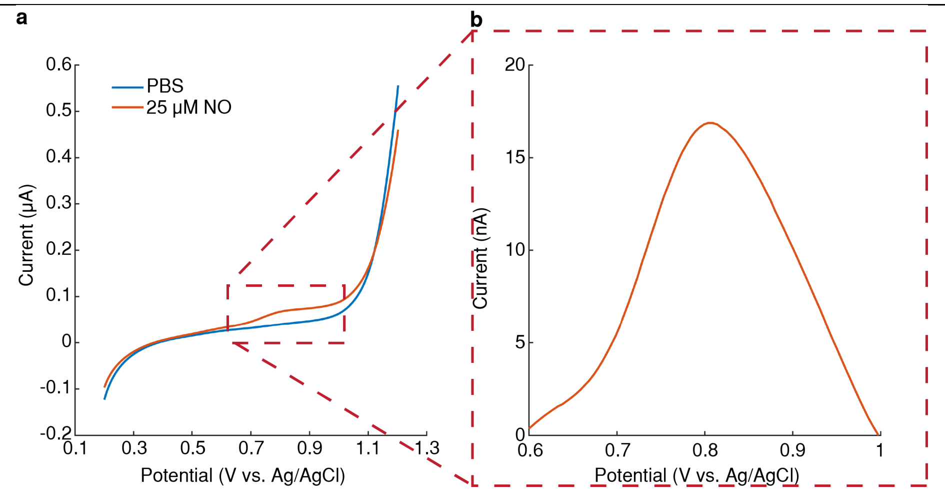


**Figure S5. Chronoamperometry (CA) operation potential characterization by linear scan voltammetry (LSV). (a)**, Determination of NO oxidation peak at 0.81V by LSV. (**b)**, Zoomed-in view of baseline subtracted LSV curve with 25 μM NO. LSV parameters: 0.2V-1.2V, 10mV/s.

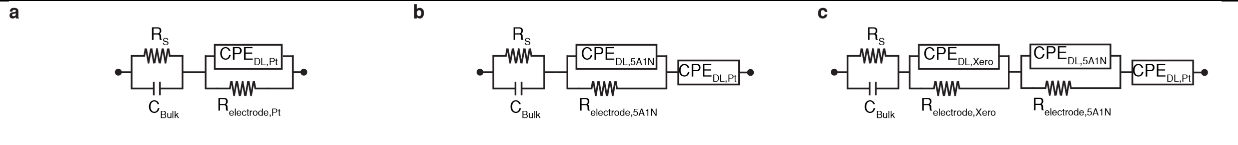


**Figure S6. Equivalent circuit model for electrode-electrolyte interfaces based on EIS data. a**, Platinum electrode. **b**, 5A1N modified platinum electrode. **c**, fluorinated xerogel-5A1N-platinum electrode.

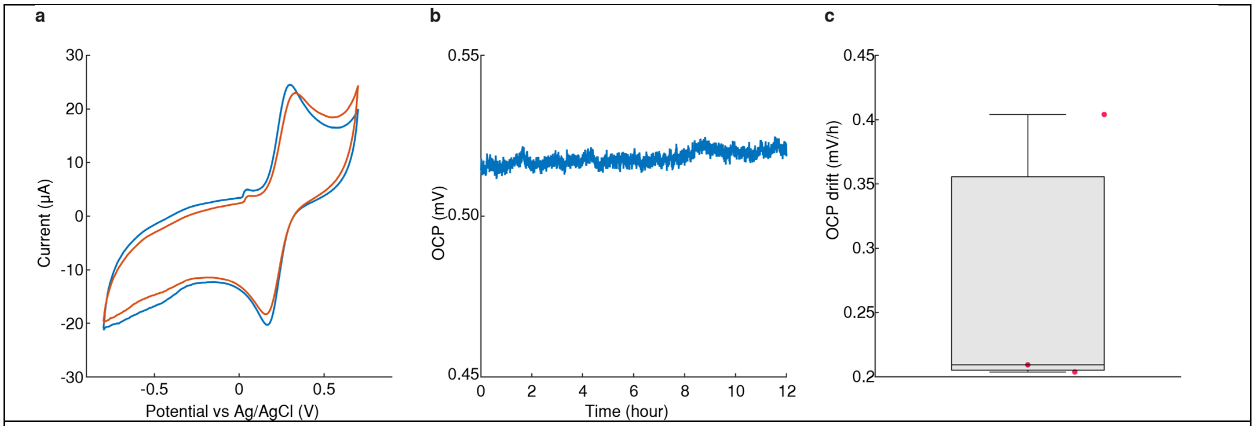


**Figure S7. On-chip reference electrode characterization. a**, Cyclic voltammetry (CV) characterization comparison by using gold disc electrode as working electrode, platinum wire as counter electrode and screen-printed Ag/AgCl ink and commercial Ag/AgCl as reference electrode in 1M $[Fe(CN)_6]^{3-}$ solution. Blue – Ag/AgCl ink as reference electrode. Orange – commercial Ag/AgCl as reference electrode. **b**, Representative long-term stability of screen-printed Ag/AgCl ink by open circuit potentiometry (OCP) measurement against commercial Ag/AgCl in 1xPBS over 12 hours. **c**, Rate of OCP drift over 12 hour period (n = 3).

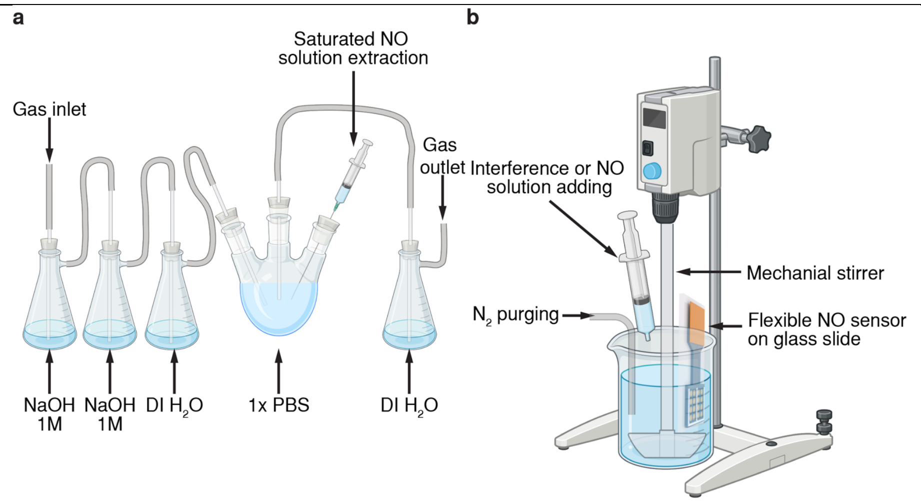


**Figure S8. MERLIN *in vitro* calibration schematic illustration. a**, NO saturated solution preparation by bubbling NO gas through 1 M NaOH, DI water and into 1xPBS in ice bath. **b**, *In vitro* setup for NO sensor sensitivity and selectivity calibration.

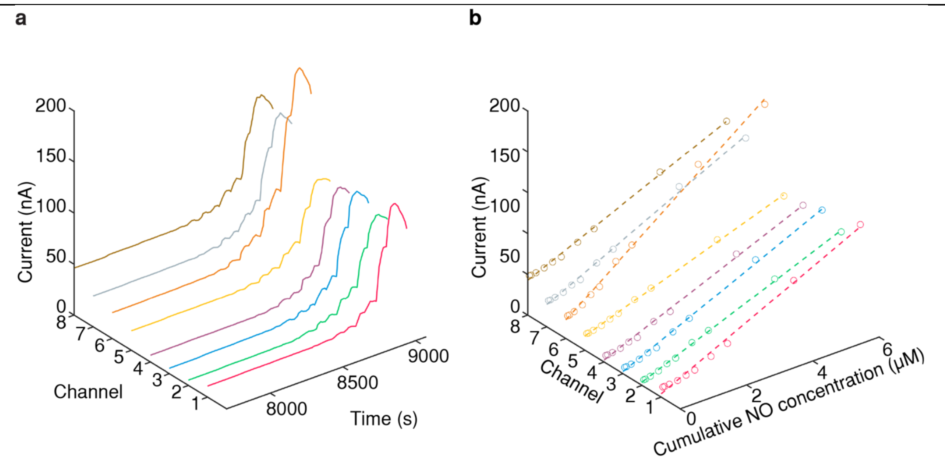


**Figure S9. Representative 8-channel multiplexed MERLIN array calibration. a,** Representative 8-channel multiplexed current measurement during calibration. **b**, Representative 8-channel calibration curve and linear regression of current against cumulative NO concentration.

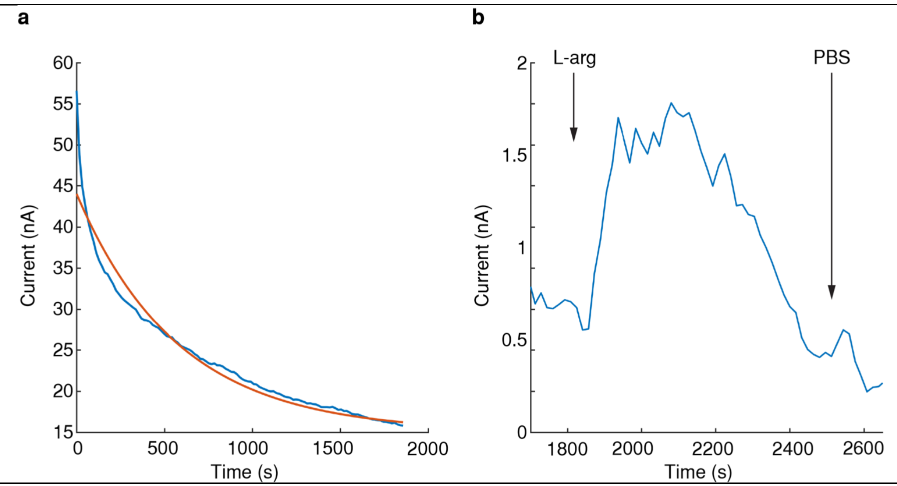


**Figure S10. Extrapolation of basal current and MERLIN function validation. a**, Representative exponential fitting to determine baseline current and baseline NO concentration. Blue – raw data collected by MERLIN. Orange – exponentially fit data. **b**, Addition of L-arg induces production of nitric oxide and validates functionality of MERLIN array. Addition of PBS was added as control.

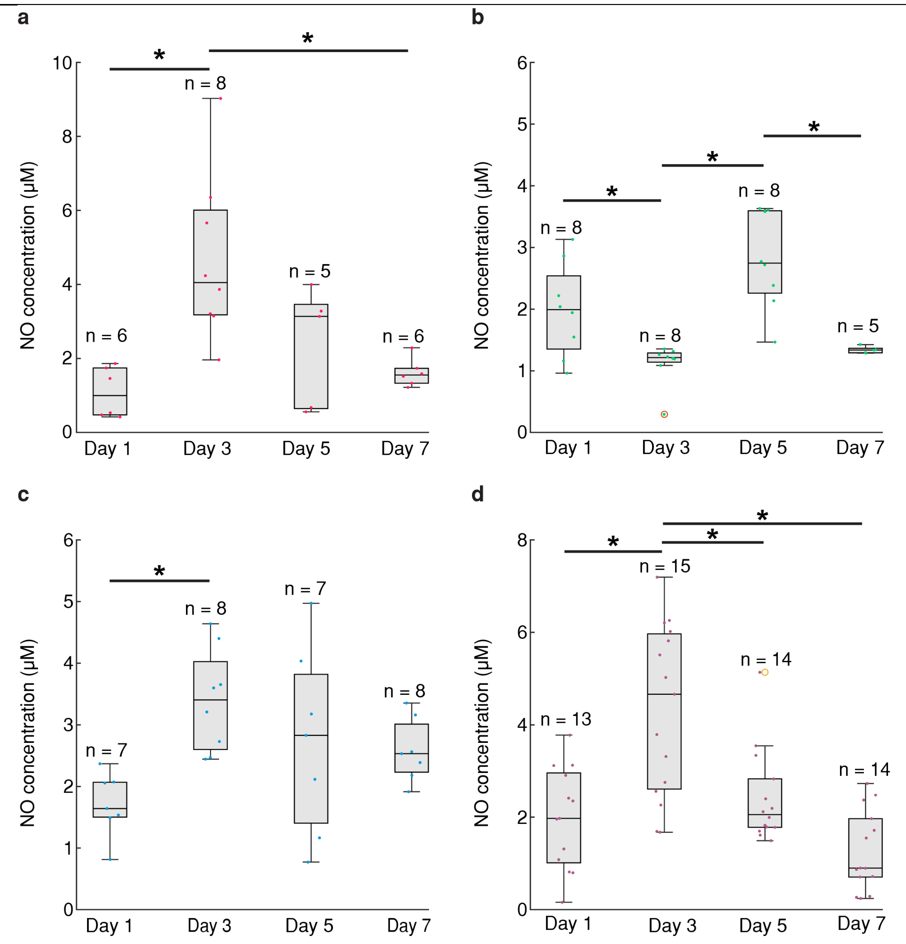


**Figure S11. Temporal NO concentration summary measured by MERLIN on 4 individual rats. a,b,c,d** Boxplot of NO concentration measured by MERLIN on rat 1, 2, 3, 4 respectively. * P< 0.05

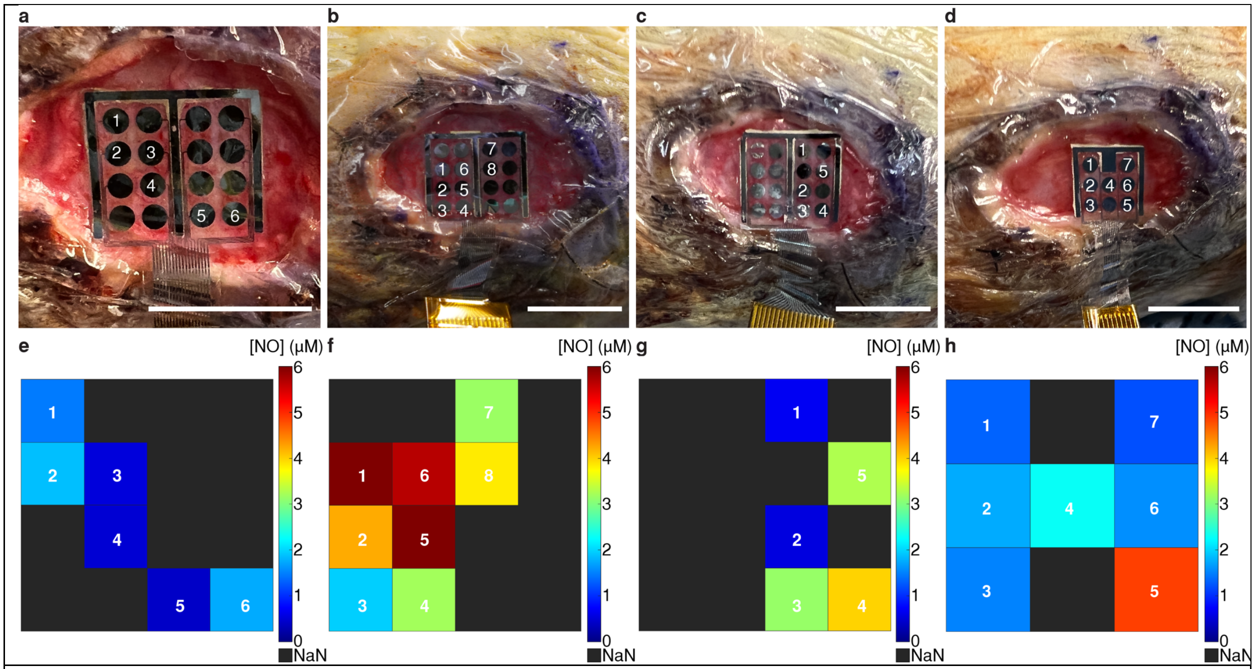


**Figure S12. Spatial NO measurement by MERLIN on rat 1 right wound. a**, **b**, **c**, **d**, MERLIN array placement on rat 1 right wound on Day1, 3, 5, and 7. Scale bar, 1cm. **e**, **f, g, h,** NO concentration mapping readout by MERLIN on Day 1, 3, 5, and 7. Color bar in units of μM NO.

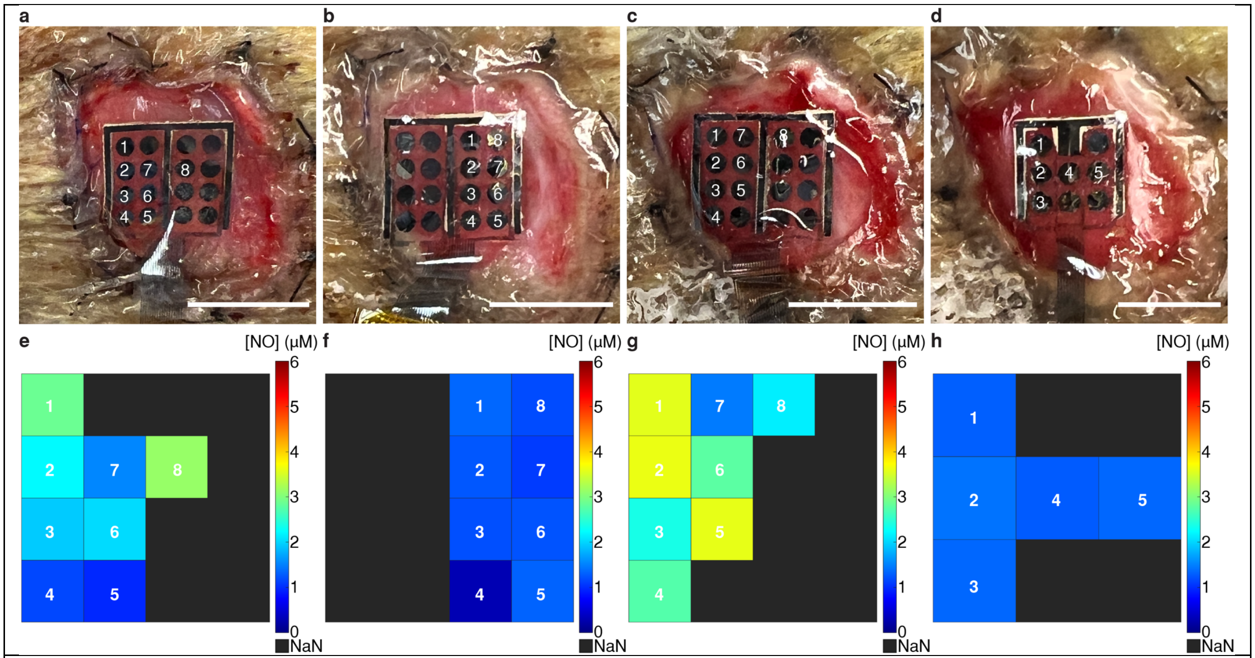


**Figure S13. Spatial NO measurement by MERLIN on rat 2 left wound. a**, **b**, **c**, **d,** MERLIN array placement on rat 2 left wound on Day1, 3, 5, and 7 post surgery. Scale bar, 1cm. **e**, **f**, **g**, **h,** NO concentration mapping readout by MERLIN on Day 1, 3, 5, and 7. Color bar in units of μM NO.

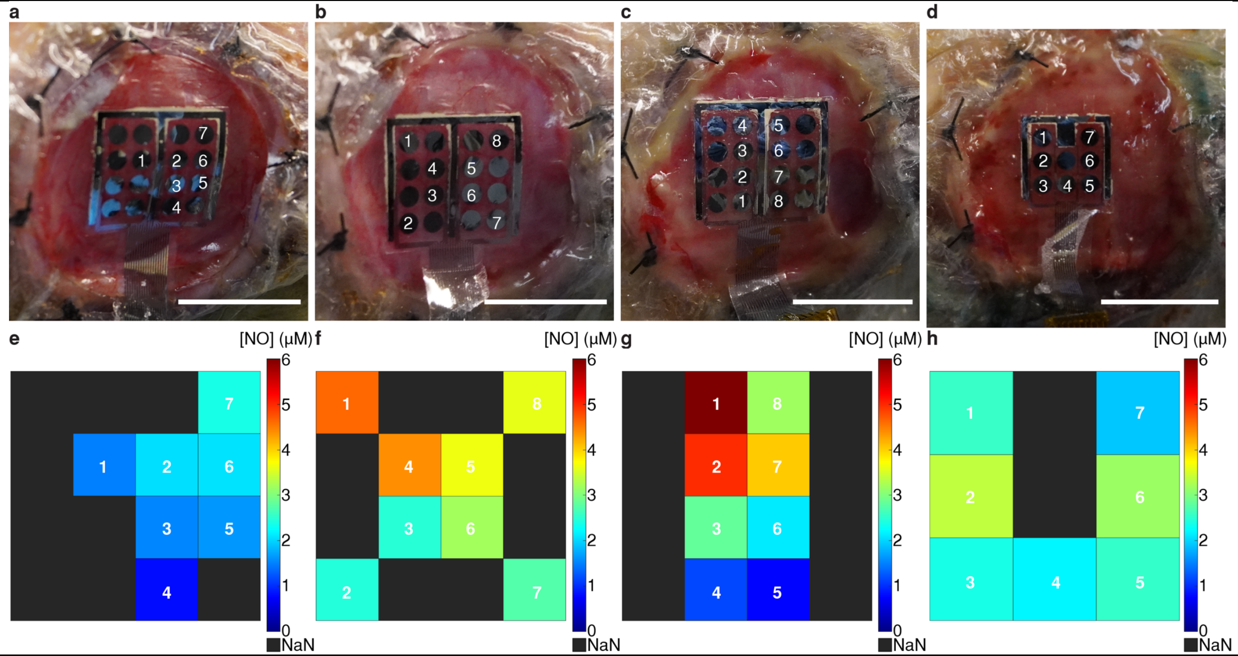


**Figure S14. Spatial NO measurement by MERLIN on rat 3 right wound. a**, **b**, **c**, **d**, MERLIN array placement on rat 3 right wound on Day1, 3, 5, and 7 post surgery. Scale bar, 1cm. **e**, **f**, **g**, **h** NO concentration mapping readout by MERLIN on Day 1, 3, 5, and 7. Color bar in units of μM NO.

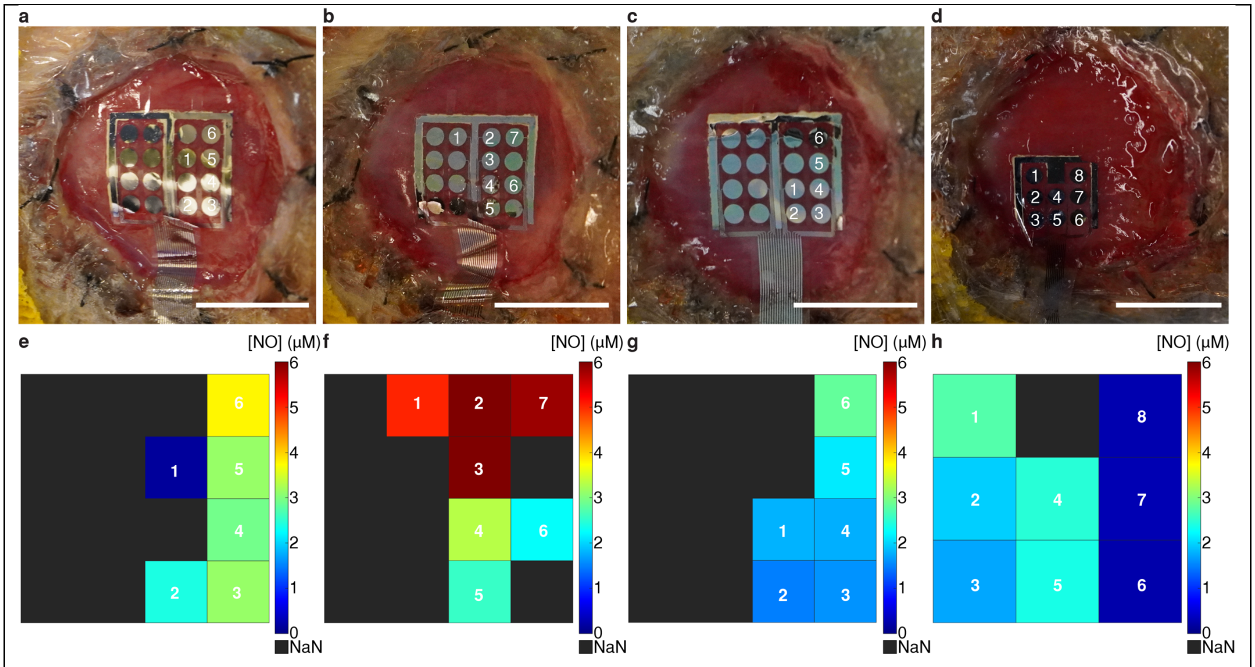


**Figure S15. Spatial NO measurement by MERLIN on rat 4 left wound. a**, **b**, **c**, **d** MERLIN array placement on rat 4 left wound on Day1, 3, 5, and 7 post surgery. Scale bar, 1cm. **e**, **f, g, h,** NO concentration mapping readout by MERLIN on Day 1, 3, 5, and 7. Color bar in units of μM NO.

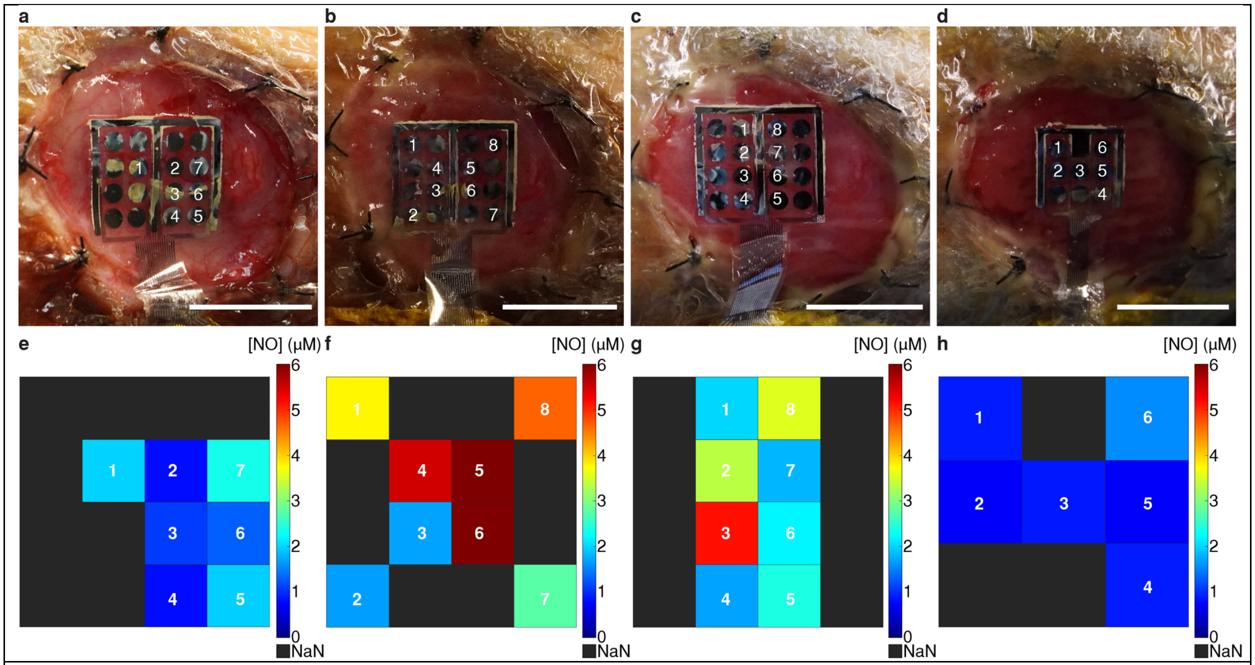


**Figure S16. Spatial NO measurement by MERLIN on rat 4 right wound. a**, **b**, **c**, MERLIN array placement on rat 4 left wound on Day1, 3, 5 post surgery. Scale bar, 1cm. **d**, **e**, **f**, NO concentration mapping readout by MERLIN on Day 1, 3, 5. Color bar in units of μM NO.

| Reference | Structure | Sensitivity per unit area | Linear Range | LOD | Selectivity | Reproducibility | in vivo measurement |
|---|---|---|---|---|---|---|---|
| Micah D. Brown and Mark H. Schoenfisch ACS Sensors 2019 4 (7), 1766-1773 | Rigid Pt-5A1N-fluorinated xerogel | 395 $nA/\mu M/cm^2$ | 0.01 - 6 μM | 1 nM | **$k\_NO_2^-$** = 4.9 | n ≥ 8 | not compatible for *in vivo* |
| Li, R., et al., *Nat Commun* 11, 3207 (2020). | Flexible Poly(eugenol)-Au -PLLA-PTMC | 26.45 $nA/\mu M/cm^2$ (0-5uM) | 10nM – 100μM | 3.92 nM | Not reported | n =3 | rabbit joint cavity *in vivo* |
| Jungmi Moon, et al., *Analytical Chemistry*, 2016 *88* (18), 8942-8948 | Rigid Platinized Pt – fluorinated xerogel | 11.7 $nA/\mu M/cm^2$ | Up to 3 μM | 9.55 nM | Not reported | n = 10 | rat brain *in vivo* |
| Ha, Y. Anal. Chem. 2016, 88 (5), 2563–2569. | Rigid Pt - Pt-black – fluorinated xerogel | 32.9 $nA/\mu M/cm^2$ | Up to 0.3uM | 6 nM | Not reported | n = 6 | Needle probe, rat brain *in vivo* |
| Santos, R. M., et al. (2013). Biosens. Bioelectron., 44, 152-159. | Rigid Hemin/Carbon nanotubes/Chitosan carbon fiber electrode | - | 0.25-1uM | 25nM | Not reported | n = 16 | Rat brain *in vivo* |
| **Our work** | **Flexible Pt-5A1N-fluorinated xerogel** | **883 ± 283 $nA/\mu M/cm^2$** | **up to 6 μM** | **8.00 nM** | **$k\_NO_2^-$ =4.44 ± 0.43 K_AA =3.58 ± 0.44 K_UA =3.84 ± 0.45** | **n = 343** | **Rat skin wound *in vivo*** |

**Table S1. Comparison among state-of-the-art NO electrochemical sensors to MERLIN.**

| | Rs (Ω) | $C_{DL,Pt}$ (S-s$^{\alpha}$) | $C_{DL,5A1N}$ (S-s$^{\alpha}$) | $C_{DL,Xero}$ (S-s$^{\alpha}$) | $X^2$ |
|---|---|---|---|---|---|
| Pt (n= 19) | $1.14 \times 10^{3} \pm 2.17 \times 10^{2}$ | $2.68 \times 10^{-6} \pm 1.41 \times 10^{-7}$ | - | - | $5.63 \times 10^{-4}$ |
| 5A1N (n= 8) | $7.97 \times 10^{2} \pm 2.40 \times 10^{2}$ | $4.67 \times 10^{-7} \pm 2.39 \times 10^{-7}$ | $5.69 \times 10^{-8} \pm 1.63 \times 10^{-8}$ | - | $7.86 \times 10^{-3}$ |
| Xerogel (n= 8) | $5.86 \times 10^{5} \pm 1.88 \times 10^{5}$ | $7.32 \times 10^{-8} \pm 4.32 \times 10^{-8}$ | $1.15 \times 10^{-8} \pm 1.73 \times 10^{-9}$ | $1.89 \times 10^{-9} \pm 8.34 \times 10^{-10}$ | $4.62 \times 10^{-3}$ |

**Table S2**. Summary of solution resistance, double layer capacitance and chi-square values of equivalent circuit fitting for EIS.

| | $\alpha_{CPE}$ in $C_{DL,Pt}$ | $\alpha_{CPE}$ in $C_{DL,5A1N}$ | $\alpha_{CPE}$ in $C_{DL,Xero}$ |
|---|---|---|---|
| Pt (n= 19) | $8.83 \times 10^{-1} \pm 5.36 \times 10^{-3}$ | - | - |
| 5A1N (n= 8) | $8.21 \times 10^{-1} \pm 3.12 \times 10^{-2}$ | $7.39 \times 10^{-1} \pm 2.70 \times 10^{-2}$ | - |
| Xerogel (n= 8) | $8.90 \times 10^{-1} \pm 3.58 \times 10^{-2}$ | $8.89 \times 10^{-1} \pm 1.33 \times 10^{-2}$ | $9.69 \times 10^{-1} \pm 2.28 \times 10^{-2}$ |

**Table S3**. Calculated constant phase element (CPE) using equivalent circuit models presented in Figure S5.
